# BioToken and BioFM – Biologically-Informed Tokenization Enables Accurate and Efficient Genomic Foundation Models

**DOI:** 10.1101/2025.03.27.645711

**Authors:** Aleksandr Medvedev, Karthik Viswanathan, Praveenkumar Kanithi, Kirill Vishniakov, Prateek Munjal, Clément Christophe, Tiago R Magalhães, Marco AF Pimentel, Ronnie Rajan, Shadab Khan

## Abstract

Genomic variation underlies human phenotypic diversity, disease susceptibility, and evolutionary adaptation. Although large-scale genomic sequencing has transformed our ability to map genetic variation, accurately modeling and interpreting this data remains a central challenge; while genomic foundation models (GFMs) are a promising approach, they suffer from fundamental limitations. Current GFMs typically treat DNA simplistically as nucleotide-only sequences, overlooking critical biological context, such as genomic annotations, regulatory elements, and functional contexts central to genomic interpretation. Here, we introduce BioToken, a modular and extendable tokenization framework designed to encode genomic variants and biologically relevant region annotations directly into genomic representations. By utilizing intrinsic inductive biases, BioToken facilitates meaningful representation learning and generalization across diverse molecular phenotypes, such as gene expression, alternative splicing, and variant pathogenicity prediction. Built on BioToken, our genomic foundation model, BioFM, achieves competitive or superior results relative to specialized models (e.g., Enformer, SpliceTransformer) and GFMs up to 7B parameters across a comprehensive suite of genomic benchmarks, including noncoding pathogenicity, expression modulation, sQTL prediction, and long-range genomic interactions. Notably, BioFM achieves state-of-the-art performance with significantly fewer parameters (265M), substantially reducing training costs and computational requirements. Our findings highlight the substantial advantages of integrating biologically-informed inductive biases into genomic foundation modeling, providing a robust and accessible path forward in genomics. We provide our code and model checkpoints to support further research in this direction.

---

Genomes encode life’s biological complexity and underpin most aspects of organisms’ function and phenotypic diversity through variations at the molecular level. With high-throughput sequencing, our ability to map and characterize genetic variants associated with diseases and phenotypic traits has improved. Despite these advances, accurately modeling and interpreting genomic data remains a challenge [1, 2]. This challenge largely arises from conceptual limitations inherent in existing modeling paradigms, specifically the treatment of DNA sequences as collections of nucleotides (A, C, G, T) without explicit consideration of their diverse biological contexts and regulatory significance within the broader genomic landscape.

Drawing inspiration from the success of Large Language Models (LLMs), recent research has adapted similar architectures and pretraining strategies to genomic data, producing Genome Foundation Models (GFMs) [3–9]. These models commonly employ tokenization and training objectives similar to those used in natural language processing, typically treating genomic sequences as strings of characters to be modeled either through sequential prediction or masked language modeling [3, 4]. While effective for generation [4, 10], this approach overlooks available biological annotations and regulatory context that are central to genome function. Unlike natural languages, where syntactic and semantic rules are relatively well-defined, genomic meaning arises from context-dependent regulatory interactions, chromatin structure, and cellular state—areas that remain subjects of active investigation [11–13]. Ignoring these inductive biases risks reducing genomes to text-like sequences without capturing their regulatory and functional organization.

In addition to overlooking regulatory annotations, GFMs must also contend with the sparsity of human genetic variation: differences between individuals typically affect less than 1% of DNA [14, 15]. This imbalance means that models trained with language-style objectives, such as autoregressive prediction or masked language modeling are incentivized to memorize common reference sequence patterns rather than learn representations that capture the functional impact of rare variants or context-dependent regulation [16, 17]. Incorporating regulatory annotations and other functional signals may provide complementary structure that helps models generalize beyond reference patterns and capture variant effects in biologically meaningful contexts. As a result, current GFMs that do not leverage such information tend to perform well on tasks where local sequence context is sufficient, such as motif discovery or promoter classification, but underperform on tasks that require generalization across diverse biological contexts, such as eQTL or variant effect prediction [18].

Recent works have highlighted these shortcomings, where they demonstrate that pretrained genomic models frequently perform no better, and sometimes worse, than randomly initialized models and supervised baselines on a range of downstream tasks [18, 19]. Such results question the utility of current genomic pretraining methods. These findings signal a clear need to revisit the assumptions of foundational modeling, developing architectures and data representations that explicitly leverage genomic structure.

To address these challenges, we introduce BioToken, a modular and extendable tokenization approach designed to incorporate genomic inductive biases by integrating biologically relevant annotations—including Single Nucleotide Variants (SNVs), insertions, and genomic features such as exons, introns, transcripts, and coding regions—directly into sequence-based genomic representation. This approach fundamentally reconceptualizes how genomic foundation models process DNA information: rather than treating sequences purely as collections of nucleotides, BioToken embeds biological annotations as specialized tokens within the sequence context, enabling models to simultaneously learn from both nucleotide patterns and their biological significance. While prior work [4] has explored specialized prompt tokens for specific adaptation tasks—such as Evo’s protein-specific conditioning tokens—BioToken integrates diverse biological context information directly during foundational pretraining rather than as task-specific conditioning labels. LoRNA-SH explored different tokenization of exons and introns [20], but BioToken uses annotations instead and is the first tokenization scheme with variant tokens. This sequence-centric approach with embedded biological context allows for the development of a single foundation model with intrinsic genomic knowledge that can be efficiently adapted across diverse downstream tasks without extensive retraining.

Importantly, BioToken is highly modular and can be easily extended to incorporate additional biological modalities such as RNA transcript variants, epigenetic markers like methylation or histone modifications, and even chromatin accessibility signals. This makes BioToken a a robust approach for different modeling tasks in the rapidly evolving landscape of genomics research. Building upon BioToken, we developed BioFM, a genomic foundation model explicitly designed for computational and parameter efficiency, utilizing recent advances in transformer architectures optimized for biological sequences. BioFM is a decoder-only transformer architecture [21], with only 265 million parameters, significantly smaller than contemporary GFMs such as Nucleotide Transformer [3] with up to 2.5B parameters, Evo1 and Evo2 [4, 10], with up to 40B parameters.

To validate BioFM and the effectiveness of BioToken, we conducted comprehensive benchmarking across diverse set of variant effect tasks, each representing distinct biological complexities. These evaluations included noncoding pathogenicity assessment (BEND benchmark [22]), gene expression modulation (eQTL dataset [23], DeepSea dataset [24]), sQTL prediction [25], coding variant pathogenicity (AlphaMissense [26]), meQTL prediction, and novel tasks such as local ancestry and rare variant classification. BioFM consistently achieved superior performance, matching or surpassing specialized models such as Enformer [27] in gene expression prediction and SpliceTransformer [28] in alternative splicing analysis, tasks previously considered challenging for generalized genomic models. Further validating our approach, BioFM demonstrated exceptional performance in other genomic tasks, outperforming existing GFMs on the Genomic Long-Range Benchmark (GLRB) [29], Trait Gym [30], and Dart-Eval [31]. Moreover, BioFM also exhibited promising generative capabilities, as demonstrated through generation of promoter sequences conditioned solely on gene inputs. This enables synthetic generative biology with potential applications in discovering novel regulatory structures given a gene as input.

Our results show that representing genomic sequences with biologically meaningful information leads to more efficient modeling, allowing BioFM to achieve strong performance while reducing model size by an order of magnitude compared to models like Evo2. By incorporating such annotations during pretraining, BioFM is not restricted to function-specific applications [27] but instead learns general features of the genome that can be adapted to a wide range of downstream tasks with minimal fine-tuning. Our research objective is to demonstrate that this biologically informed pretraining strategy provides a practical framework for genomic foundation models, linking sequence information with existing biological knowledge. In the longer term, such models could support deeper studies of molecular phenotypes, disease mechanisms, and applications in personalized medicine [32].

## Results

### Biologically-Informed Tokenization of Genomes with BioToken

We developed BioToken, a modular tokenization framework design that encodes variants and biological annotations directly within DNA sequences. Conventional genomic modeling simplifies DNA sequences as strings of nucleotides, ignoring biological context essential for accurate genomic interpretation. In contrast, to help the model focus on genetic variations, we developed a tokenization strategy that replaces standard nucleotide tokens (A, G, C, T) with special variant tokens (Ă, Ğ, Č, Ť) to capture single nucleotide variants (SNV) and insertions. Similarly, BioToken introduces special tokens that encode genomic annotations such as transcript boundaries, exon-intron architecture, and coding DNA sequences (CDS) (Appendix A.2.1) (Figure 1a). Encoding these biologically meaningful annotations inherently leverages genomic inductive biases, shifting the learning objective from superficial positional memorization towards deeper functional representation learning. Notably, the modularity and extendibility of BioToken enables it to integrate additional genomic annotations not explored in this study, such as RNA secondary structures, epigenetic states (e.g., methylation, histone modifications), and chromatin accessibility patterns. BioToken therefore provides a foundation for genome-scale modeling across diverse downstream applications.

**Fig. 1:**
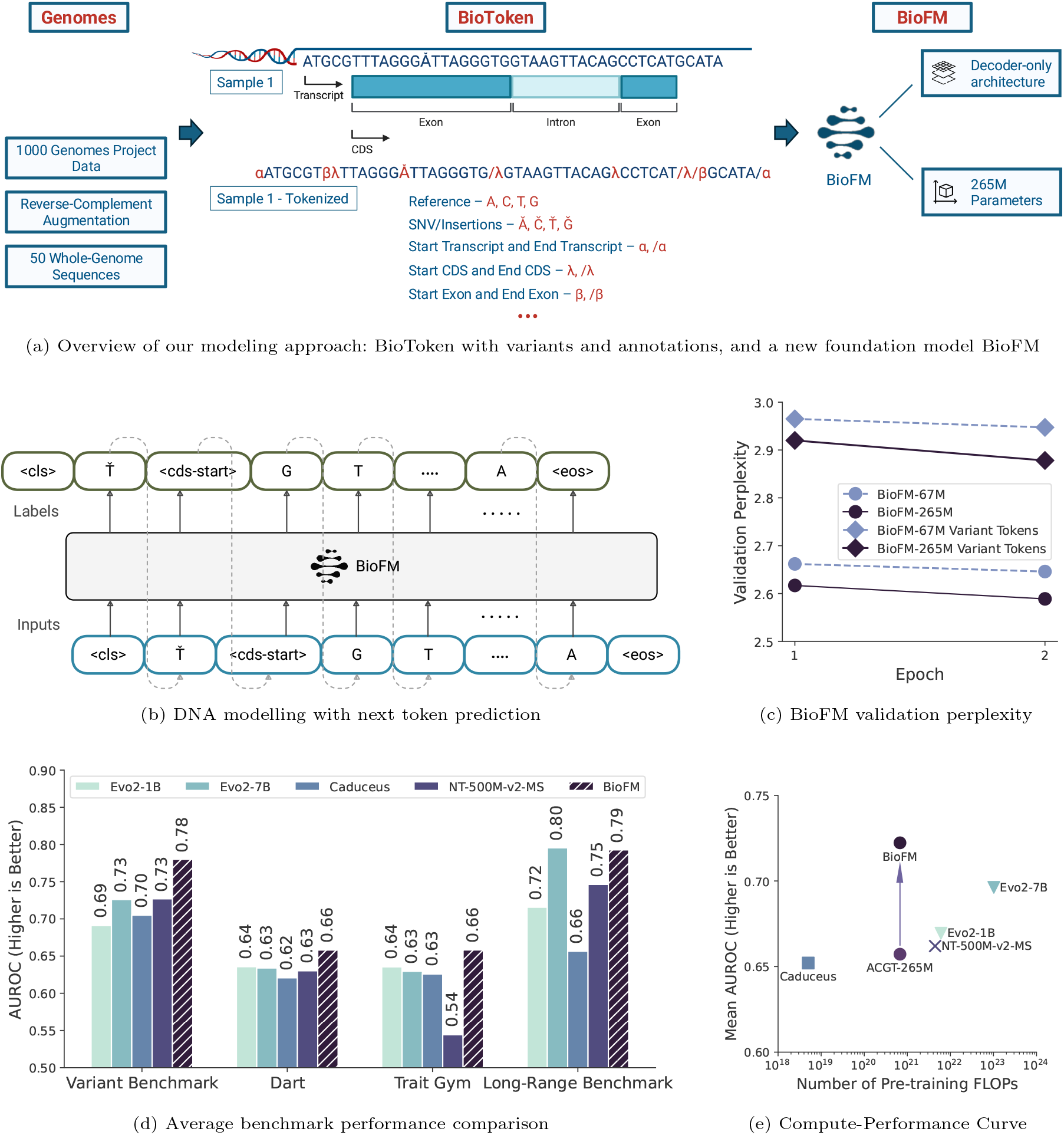
BioFM integrates biologically-informed tokenization, significantly outperforming GFMs and specialized models across a range of tasks and benchmarks. **a)** Schematic representation of BioFM’s architecture highlighting its key components and an illustrative example showing how DNA sequences are annotated. **b)** BioFM employs next token prediction to model standard nucleotides, variants and annotations. **c)** BioFM validation perplexity decreases with increasing size and training tokens for variant and other tokens. **d)** BioFM outperforms every GFM up to 7B on three out of four variant effect benchmarks. **e)** Adding annotations to the nucleotide-only model with the same size and amount of compute (ACGT-265M) significantly increases variant effects performance across four benchmarks from (d).

### BioFM – Computationally Efficient Foundation Modeling

Using BioToken, we developed BioFM, a computationally efficient genome foundation model optimized for biologically-informed representation learning. BioFM employs a decoder-only transformer architecture comprising 265M parameters (Figure 1b), substantially smaller than contemporary GFMs, which significantly reduces computational resource demands. To systematically exploit DNA’s reverse-complement symmetry, we implemented reverse-complement augmentation during training, randomly applying it to 50% of input sequences while applying only strand-specific annotations. This approach compels the model to learn symmetrical representations, which is essential given that many genomic regulatory and functional motifs exhibit strand symmetry. BioFM was pretrained on WGS data from 50 individuals (Table A1) selected from all five superpopulations from 1000 Genomes project [14], yielding approximately 166 billion tokens for one epoch of pretraining (we trained BioFM for two epochs). After evaluating multiple context lengths, we empirically determined 6K nucleotide as the ideal context window. Further, we adopted Rotary Position Embeddings (RoPE) but adjusted the positional embedding frequency parameter *θ* from 10^4^ to 10^5^, to mitigate positional biases and effectively model genomic dependencies over longer distances [33]. Our initial ablation analyses was conducted using a reduced-size 67M-parameter model, and all subsequent experiments were perfomed using a scaled-up 265M-parameter model.

### BioFM Accurately Predicts Variant Effects Across Diverse Molecular Phenotypes

We developed a new Variant Benchmark to evaluate BioFM’s ability to effectively utilize variant information across multiple biological contexts, significantly extending beyond conventional genomic benchmarks [3, 22, 29, 34, 35], which primarily emphasize region classification tasks. Although existing assessments such as BEND [22] and the Genomic Long-Range Benchmark (GLRB) [29] measure critical functions like noncoding pathogenicity and tissue-specific expression, they do not represent the broader spectrum of variant-mediated molecular processes. In contrast, we organize our benchmark evaluations into two major categories to systematically assess BioFM’s capabilities: **(1)** Functional Effects of Genetic Variants, focusing on the direct molecular and biological consequences of genetic variation, and **(2)** Population-Level Variation, assessing BioFM’s sensitivity to broader genetic patterns across human populations.

#### 1. Functional Effects of Genetic Variants

We assessed BioFM’s capability to model variant effects across five key molecular processes.

- **Coding pathogenicity assessment**: Accurate prediction of pathogenic coding variants is fundamental to precision medicine and clinical genomics. We assessed this using the AlphaMissense dataset [26], comprising a comprehensive catalog of coding variants annotated for pathogenicity.
- **Noncoding pathogenicity assessment**: Pathogenic variants in noncoding regions significantly impact gene regulation, influencing many complex traits and diseases. To evaluate predictive accuracy, we employed the BEND dataset, containing 295,000 annotated SNVs in noncoding genomic regions [22].
- **Expression effect prediction**: Variant-driven changes in gene expression contribute to phenotypic diversity and disease processes. To test BioFM’s representation of these regulatory effects, we used the eQTL dataset from Borzoi [23] and expression dataset from DeepSea [24].
- **Alternative splicing**: Variant-induced alternative splicing contributes significantly to human proteomic diversity and affects biological processes and diseases. To evaluate modeling accuracy for splicing variations across multiple cellular contexts, we made use of sQTL dataset from Borzoi [23].
- **DNA methylation**: Variant-driven changes in DNA methylation patterns modulate gene-environment interactions and influence diverse developmental and disease phenotypes. To test the model’s ability to predict variant effects on DNA methylation, we utilized methylation quantitative trait loci (meQTL) data from the GRASP database [36].

#### 2. Population-Level Variation

We next evaluated BioFM’s ability to capture population-level insights. Specifically, we considered two tasks:

- **Local ancestry classification**: Understanding how genetic variants collectively encode ancestral backgrounds informs studies of population structure, evolutionary biology, and disease susceptibility. From the mechanistic perspective, local ancestry classification tests sensitiviy of GFM representations to small changes in long sequences. To evaluate this capability, we used genomic segments labeled by five major superpopulations from the 1000 Genomes Project [14].
- **Common vs synthetic variants**: This task evaluates the model’s ability to recognize biologically conserved genomic contexts characteristic of authentic common variants. We randomly sampled 100K common variants (MAF *>* 0.05) from GnomAD [15] and paired each with a synthetic control (“rare”) variant generated by randomly substituting a nucleotide within a ±16-nucleotide local context window.

#### Benchmarking Protocol and Model Evaluation

We developed a standardized assessment protocol to fairly evaluate different model architectures across all tasks. For each variant, we use its DNA context up to the maximum length supported by the model, capped at 12K nucleotides. This limit was set because only HyenaDNA and Caduceus can process longer sequences, but their performance plateaus beyond this context length. Our evaluation approach differs slightly between encoder and decoder-only architectures to exploit their respective strengths. For encoder models, we center the variant within its context, using equal length of sequence information from both upstream and downstream regions. For decoder-only models, namely BioFM and HyenaDNA, we extract embeddings from both upstream and downstream sequences of equal length (half the evaluation context size each), with the downstream sequence reverse complemented. We generate separate embeddings for reference and mutated sequences, average the upstream and downstream embeddings for each, then concatenate the averaged reference and mutated vectors to create the final variant representation. Refer Section A.4 for complete implementation details. The final variant embeddings serve as inputs to a linear classifier. We avoid more complex nonlinear methods or full fine-tuning, as these could obscure differences in embedding quality and require extensive hyperparameter tuning for each model.

We evaluated six GFMs—Evo2-7B-Base, Evo2-1B-Base, NT-500M-v2-MS, NT-500M-1000G, HyenaDNA, Caduceus—on the Variant Benchmark. We selected models with different sizes, amount of compute spent ranging from 100x to 0.01x of BioFM ( Figure 1e), context length from 6144 to 1M tokens. Performance for all of them is shown in Figure 2a. Additionally, we included GENA-LM, DNABERTv2, and a supervised convolutional neural network (CNN) with 120K parameters as a baseline, trained separately for each task and fold. Results for all models are shown in Figure A2.

**Fig. 2:**
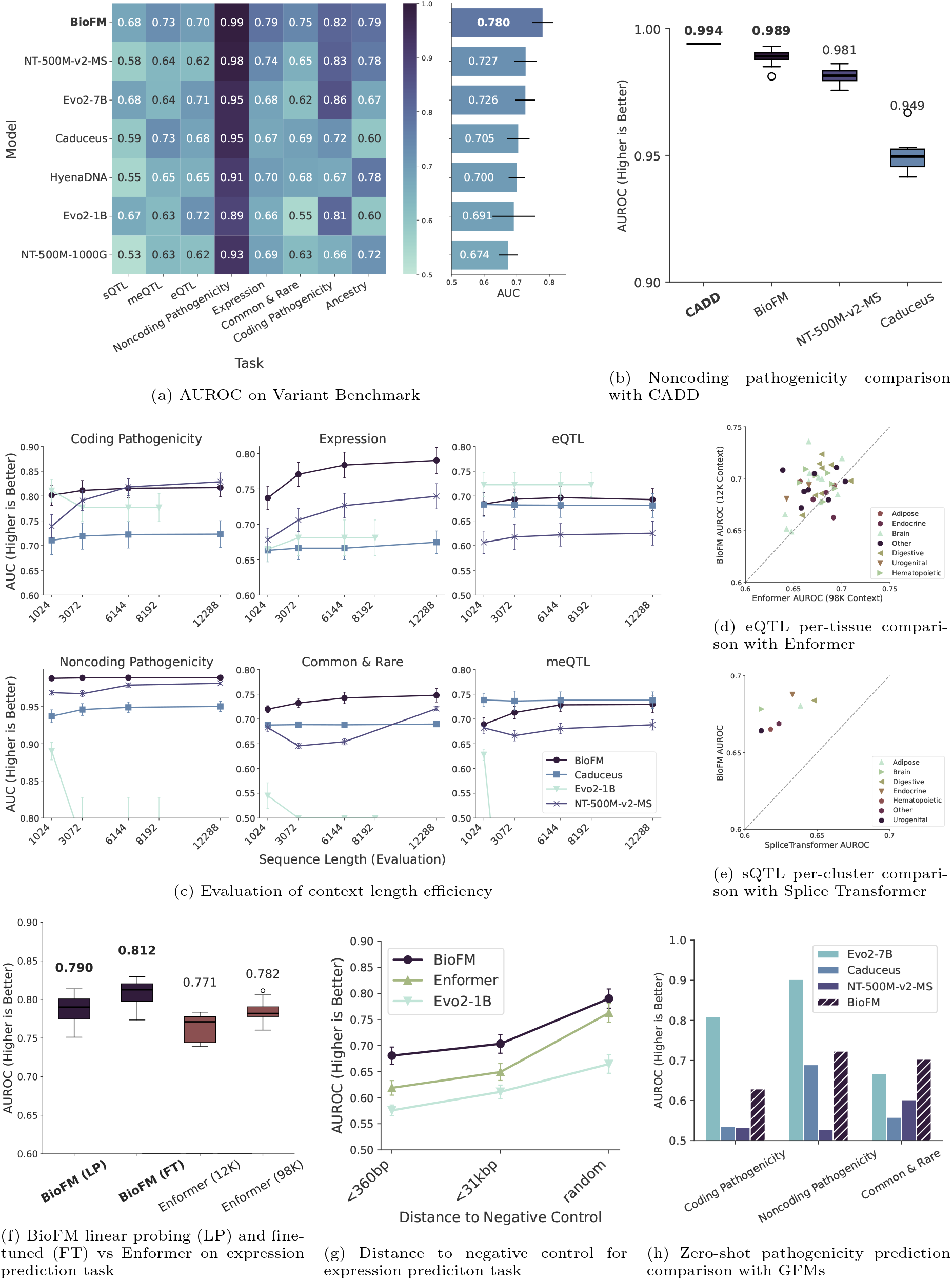
Extensive evaluation shows that BioFM does well in modeling variants, and efficiently using genomic context for superior accuracy and generalization. Comprehensive performance evaluation of BioFM across multiple genomic prediction tasks using 11-fold crossvalidation. **a)** Across a broad spectrum of variant prediction tasks, BioFM outperforms other GFMs. Error bars are 2 standard deviations wide, centered on the mean. **b)** BioFM shows comparable performance to CADD for noncoding pathogenicity using linear probing finetuning strategy. **c)** BioFM demonstrates robust scalability and accuracy across increasing context lengths, consistently outperforming competing models, particularly in noncoding pathogenicity, rare variant, and expression prediction tasks. Error bars are 2 standard deviations wide, centered on the mean. **d)** BioFM achieves consistent, tissue-independent performance improvements over SpliceTransformer on tissue-specific splicing (sQTL) predictions. **e)** BioFM significantly outperforms Enformer on a tissue-specific eQTL dataset. **f)** BioFM with linear probing and finetuned versions both improves expression predictions, exceeding Enformer performance even when Enformer utilizes significantly longer context windows. Box plot shows median, 1st and 3rd quartile, and outliers. **g)** AUROC for BioFM, Enformer, and Evo2-1B on expression prediction using negative controls restricted to *<*360 bp, *<*31 kbp, and randomly selected variants. BioFM consistently achieves higher performance across all settings, indicating robustness to distance-based control selection. **h)** BioFM demonstrates competitive zero-shot variant pathogenicity prediction capabilities compared to other large-scale genomic foundation models, falling only behind Evo2-7B on 2 out of 3 tasks.

BioFM consistently generated superior representations for variant effects prediction (overall *AUROC* = 0.780) relative to second-best NT-500M-v2-MS (*AUROC* = 0.727), despite nearly half the parameter count Figure 2a. BioFM was able to outperform Evo2-7B which used approximately 100x more compute during pretraining on five out of eight tasks. Notably, BioFM significantly outperformed NT-500M-1000G, highlighting limitations of fixed-length k-mer tokenization for variant-rich data. BioFM achieved the highest performance across six of eight tasks, with significant advantages particularly in distinguishing common versus rare variants (*AUROC* = 0.746 vs. 0.69) and expression prediction (*AUROC* = 0.786 vs. 0.74). Importantly, we do not claim that BioFM outperforms CADD, AlphaMissense, or GPN-MSA on pathogenicity prediction Figure 2b tasks, however no GFM was able to do that in our evaluations.

#### Context Length Utilization and Embedding Quality

In general, transformer based GFMs are known to struggle to fully utilize their context windows [37]. To investigate how feature extraction is affected by context length, we evaluated model performance using variant contexts ranging from 1K to 12K nucleotides (Figure 2c).

BioFM demonstrates strong performance even with a 1K-nucleotide context, significantly outperforming NT-500M-v2-MS across all tasks, including coding pathogenicity. However, NT-500M-v2-MS scales slightly better for pathogenicity tasks. For expression and rare variants prediction, BioFM’s performance does not saturate, suggesting potential benefits from models with even longer context windows. BioFM effectively utilizes its full 6K-nucleotide window (6K in forward strand and 6K in reverse strand). By contrast, Caduceus, despite being designed for long contexts, plateaued around 3K nucleotides across all tasks. Based on these findings, we limited evaluation contexts to a maximum of 12K nucleotides. Surprisingly, the performance of larger models such as Evo2-1B and Evo2-7B decays after 1K nucleotides for four out of six tasks. This is likely due to their mean pooling strategy for extracting features. The detailed discussion of Evo2 evaluation protocol is in Methods. Across all context lengths, BioFM was the only model consistently ranking among the top two performers.

#### Direct Comparison with Specialized Genomic Models

Next, we compared BioFM to strong supervised baselines: Enformer for expression and tissue-specific eQTL prediction and Splice Transformer for sQTL prediction. Since both models generate tissueor cell type-specific predictions, and their prediction tracks do not always match with tracks in our data, we extracted their sequence embeddings for downstream linear probing to ensure a fair comparison. As shown in Figure 2f, BioFM matches Enformer’s performance on expression prediction when both models use 12K context, making it the first ever GFM to do so. Next, we finetuned BioFM with parameter-efficient LoRA on expression prediction dataset and compared it with Enformer using 98K context (Figure 2f). BioFM significantly outperformed Enformer, median *AUROC* = 0.812 vs *AUROC* = 0.782 (*p* = 0.002 subsection A.5). BioFM with linear probing also outperforms Enformer on eQTL dataset from Borzoi [23], *AUROC* = 0.697 vs *AUROC* = 0.674 (Figure 2d). Moreover, BioFM is significantly better than Splice Transformer for sQTL prediction (Figure 2e), achieving median tissue *AUROC* = 0.676 compared to *AUROC* = 0.629 of Splice Transformer. Analyzing tissue-cluster-specific performance (Figure 2e) reveals that BioFM consistently performs better across all tissue clusters. However, BioFM is not able to outperform CADD on noncoding pathogenicity prediction, albeit it is better than other GFMs (Figure 2b).

#### Computational Efficiency and Generalization Compared to Large Models

For many models, the likelihood of a nucleotide at a given position is correlated with its pathogenicity. To evaluate the zero-shot performance of BioFM, we compared it with Evo2-7B [10], which was trained on 3T nucleotide tokens spanning all domains of life. We restricted the context size to 1K nucleotides, as Evo2 models exhibit reduced performance at longer sequences, a trend observed in our experiments (Figure 2c) and also reported in the original Evo2 paper [10].

On pathogenicity prediction tasks, Evo2-7B achieved higher accuracy, but it performed worse on the Common & Rare classification task. In contrast, BioFM produced competitive results while being approximately two orders of magnitude more compute-efficient in generating variant representations (Figure 2a). This efficiency stems from its use of variant tokens and annotations, which allow the model to encode variation explicitly rather than relying solely on scale. The performance boost on the Common & Rare task can be attributed to both the incorporation of variant tokens, as demonstrated in our ablation studies (Figure 5a), and training on 1000G data. Importantly, this task is particularly challenging because common and rare variants were deliberately chosen to be in close proximity within small genomic regions, requiring models to be sensitive to subtle, local nucleotide frequency shifts.

Further, authors of NT and Evo2 [3, 10] have shown that optimal performance on downstream tasks typically requires selecting variant representations from intermediate layers, a process involving computationally expensive layer-wise evaluations. To determine whether BioFM similarly requires intermediate-layer selection, we compared the performance of variant representations from BioFM’s intermediate layers to those from its last layer (Figure A1). Our analysis demonstrates that BioFM’s last-layer embeddings are consistently superior or equivalent to intermediate-layer embeddings across tasks. Thus, last-layer embeddings in BioFM serve as universally effective representations for downstream applications, avoiding the computational overhead of per-task layer selection.

In conclusion, these findings establish BioFM as a meaningful advance in genomic modeling it is currently the only genomic foundation model that demonstrates practical utility for both variant expression and splicing effects, matching or exceeding the performance of specialized models designed specifically for these tasks.

### BioFM Performs Well on GLRB, Dart, Trait Gym Benchmarks

We evaluated BioFM on three additional benchmarks: the Genomic Long-Range Benchmark (GLRB) [29], Dart-Eval [31], and Trait Gym [30]. GLRB includes noncoding pathogenicity, expression, and OMIM-versus-common variant prediction tasks, though its datasets were preprocessed and filtered differently from those in our Variant Benchmark. Dart focuses on dsQTLs in African and caQTLs in Yoruba populations, representing molecular phenotypes not covered in Variant Benchmark. Trait Gym assesses the detection of variants influencing Mendelian and complex traits, simulating GWAS-style inference.

BioFM outperforms NT-500M-v2-MS and other models on GLRB (*AUROC* = 0.793), despite NT-500M-v2-MS being finetuned for extended context lengths (12K to 66K nucleotides, Figure 3a). This suggests that inductive biases introduced through BioToken are more effective than finetuning or context extension alone, and that GLRB tasks may not strongly depend on long-range interactions. On Dart, BioFM performs significantly better on Yoruba caQTLs, though not on the more diverse African dsQTL dataset (Figure 3c), likely reflecting both the explicit use of variant tokens and the presence of two Yoruba samples in pretraining (NA19258, NA18930). On TraitGym, BioFM shows strong performance in identifying variants underlying Mendelian traits (Figure 3d), likely reflecting the model’s ability to leverage biological annotations and variant-aware tokenization.

**Fig. 3:**
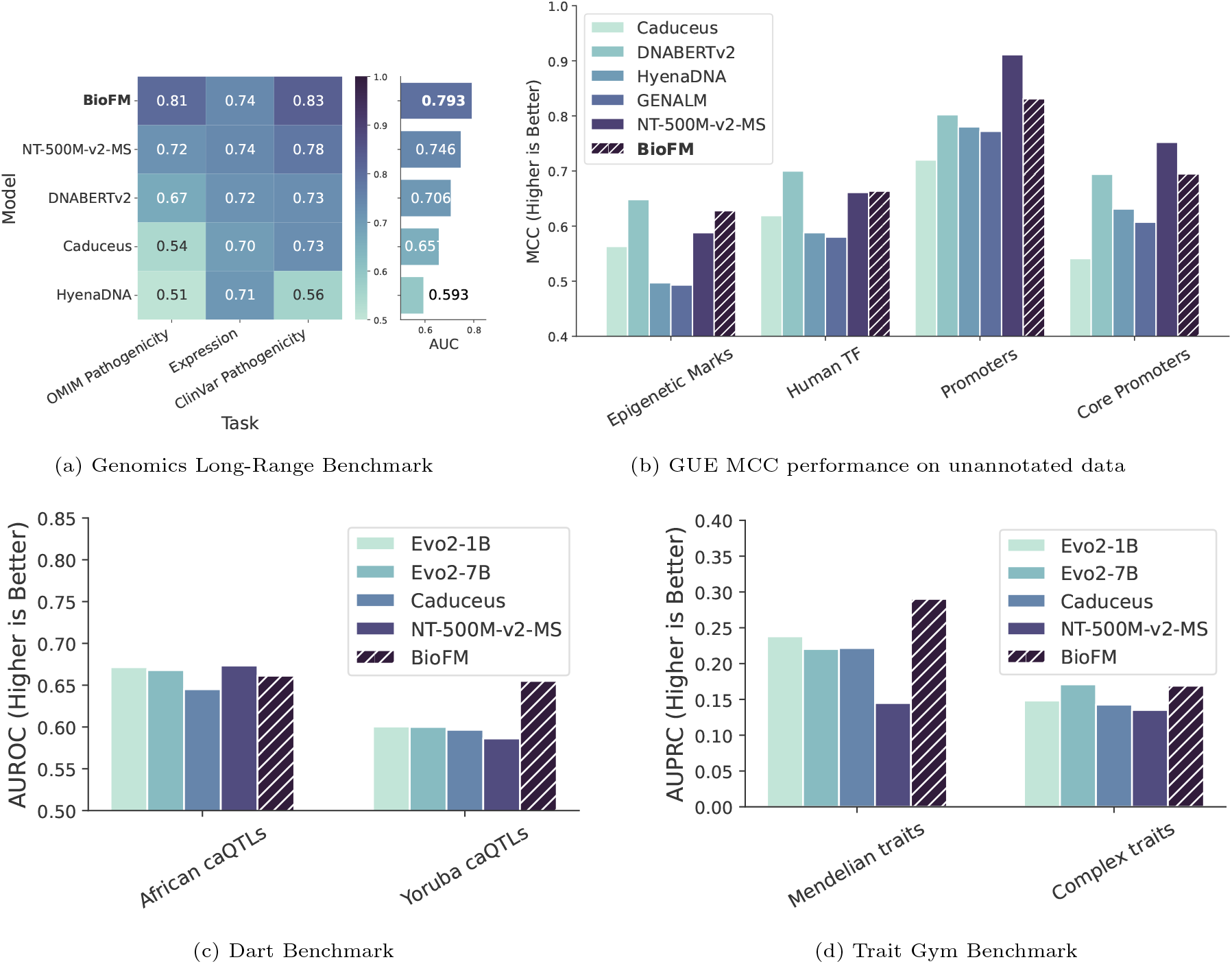
BioFM substantially improves predictive accuracy on genomic long-range benchmarks (GLRB). Without access to BioToken, in an evaluation of worst-case performance, it achieves competitive results across genomic functional annotations in the Genomic Understanding Evaluation (GUE). **a)** On GLRB, BioFM outperforms previous GFMs that required extensive fine-tuning [29] and longer genomic contexts, demonstrating more efficient usage of shortand medium-size context. We used the same test set as [29]. Results for all models except BioFM were obtained from [29], BioFM was evaluated with linear probing, other models were finetuned. **b)** Comprehensive performance evaluation on the GUE benchmark reveals BioFM consistently excels in functional genomic annotations even when specialized tokens are excluded, highlighting its generalization ability and understanding of of intrinsic genomic regulatory features. Every model underwent full finetuning with the same hyperparameter tuning budget. Detailed performance metrics are provided in Appendix, Table A5. BioFM outperforms all models on Yoruba caQTLs and is competitive on African caQTLs. We used linear probing for every model, context length is 12K except Evo2-1B and Evo2-7B, where we set it to 1K. **d)** BioFM outperforms all models on Mendelian traits task and is second-best on Complex trats task. Metric is AUPRC, following [30]. We performed all evaluations on our codebase, matching the scheme from [30] as closely as possible.

Overall, across GLRB, Dart, and Trait Gym, BioFM outperforms both longer-context models such as Caduceus and substantially larger models such as Evo2-7B, underscoring the value of variant-aware tokenization and genomic pretraining.

### BioFM Captures Broad Functional Genomic Signals

To assess BioFM’s general applicability as a foundation model, especially in scenarios where annotations are not available (and therefore annotation tokens are not included as part of the input), we evaluated its performance on the Genomic Understanding Evaluation (GUE) benchmark [34]. GUE benchmark encompasses human-centric genomic tasks, including prediction of epigenetic modifications, transcription factor binding sites, and promoter region characteristics, structured as binary region classification problems analogous to those in the NT benchmark [3]. Notably, these evaluations were conducted without BioFM’s specialized features (BioToken’s annotation tokens, variant tokens were not used, reverse complement augmentations was omitted), resulting in a lower-bound result for BioFM.

BioFM demonstrated consistently strong performance across all task categories within the GUE benchmark, ranking second overall. It was surpassed only by DNABERTv2 on epigenetic mark and transcription factor binding predictions, and by NT-500M-v2-MS on promoter-specific tasks. Nonetheless, BioFM consistently outperformed competing models such as HyenaDNA, Caduceus, and GENA-LM across all evaluated categories. Further, despite having nearly half the parameter count and lacking domain-specific adaptations, BioFM surpassed NT-500M-v2-MS on epigenetic mark and transcription factor binding tasks. These results demonstrate that BioFM remains a useful model even in applications where it doesn’t fully utilize its strengths.

### Instruction-Tuning Enables BioFM to Generate Regulatory Sequences

The generation of synthetic promoters represents a strong test for evaluating BioFM’ generative capabilities. The ability to design synthetic promoters can enable precise control over transcriptional activity, with applications in optimizing gene expression for research and therapeutic interventions [38, 39]. As promoters are typically located upstream of the corresponding gene sequence, BioFM requires fine-tuning to generate them effectively. Applying the ideas of instruction tuning used in training language models for this task, we prompt model with a gene sequence and a special token ^*′*^*P* ^*′*^ after it, creating instruction set of `⟨DNA sequence⟩ ⟨Task-identifier token⟩ ⟨Output⟩` . Training was done using instruction-answer pair of genes-promoters set and testing on a held-out set. Promoter sequences were drawn from the EPDNew dataset [40]

We systematically validated BioFM-generated promoters using three independent biological criteria: (1) compositional similarity to natural promoter sequences, and (2) accurate modeling of biologically significant promoter types (TATA versus non-TATA).

First, we assessed compositional similarity by analyzing k-mer distributions of generated promoter sequences. We found that after 10 epochs of fine-tuning, BioFM-generated promoters successfully captured the k-mer distributions of wildtype promoters, indicating convergence toward biologically realistic promoter compositions (Figure 4b).

**Fig. 4:**
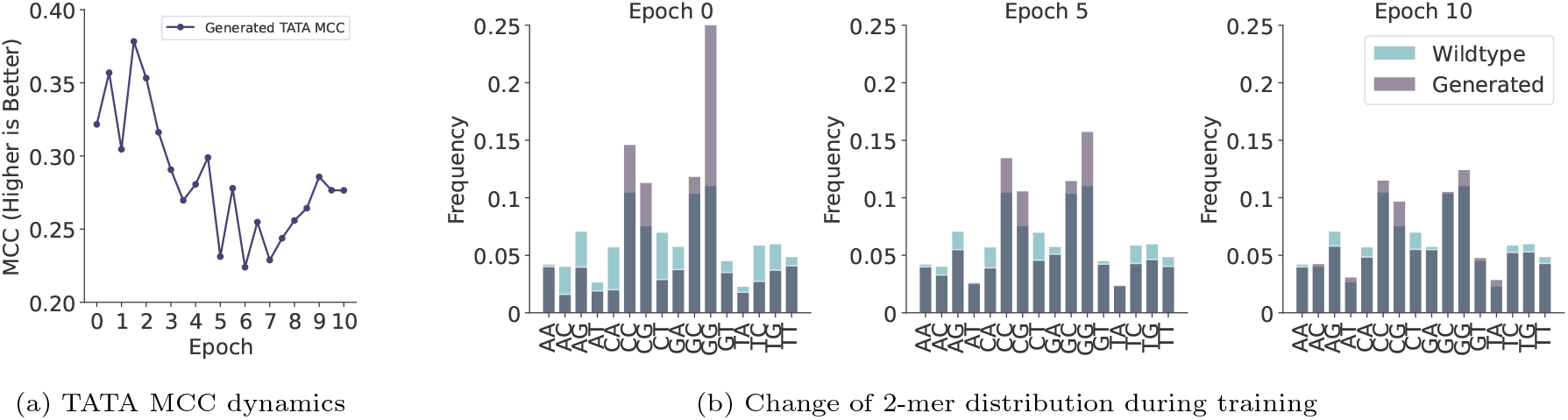
BioFM effectively generates biologically plausible synthetic promoter sequences, progressively aligning with naturally occurring promoter features through fine-tuning. **a)** MCC tracking of TATA prediction accuracy across 10 training epochs. Performance is evaluated based on the model’s ability to correctly generate TATA-box-containing promoters for genes naturally possessing wildtype TATAbox promoters (true positives) while avoiding incorrect TATA-box insertions in naturally non-TATA promoters (false positives). **b)** Comparative analysis of 2-mer frequency distributions between generated and wildtype promoter sequences, demonstrating progressive convergence of synthetic promoter sequence composition toward natural distributions during the finetuning process.

Second, we measured BioFM’s ability to model biologically significant promoter types accurately. Notably, the Matthews Correlation Coefficient (MCC) for predicting TATA-containing promoters was 0.28 after fine-tuning (Figure 4a), confirming the model’s capability in discriminating genes more likely to possess TATA promoters. Interestingly, we observed an early training phenomenon where the TATA MCC peaked higher (approximately 0.38 within the first two epochs), coinciding with the generation of promoter sequences enriched in CC and GG dinucleotides compared to the wildtype distribution. This suggests an initial overfitting to easily identifiable motifs, which was subsequently refined toward wildtype distribution.

Collectively, these evaluations indicate that BioFM effectively internalizes the relationships between genes and promoters, applying this understanding to generate biologically plausible synthetic promoters for previously unseen genes.

### Ablation, Scaling, Representation Analysis, and Attention Specialization

We assessed the contributions of BioToken and genomic variance data to BioFM’s performance by performing systematic ablation experiments, embedding analyses, and evaluations of sample efficiency.

First, we quantified the impact of BioToken and genetic diversity through ablation experiments, where we ablated the variant and annotation tokens (Figure 5a). We established baseline comparisons by training a nucleotide-only model (ACGT-265M) on the same subset from 1000 Genomes data, without any specialized tokens. Then, we assessed the specific impact of annotation tokens by training an intermediate model that included variant tokens but excluded annotations. Additionally, during evaluation, we selectively removed annotation and/or variant tokens from test sequences to quantify their effect on model’s performance.

**Fig. 5:**
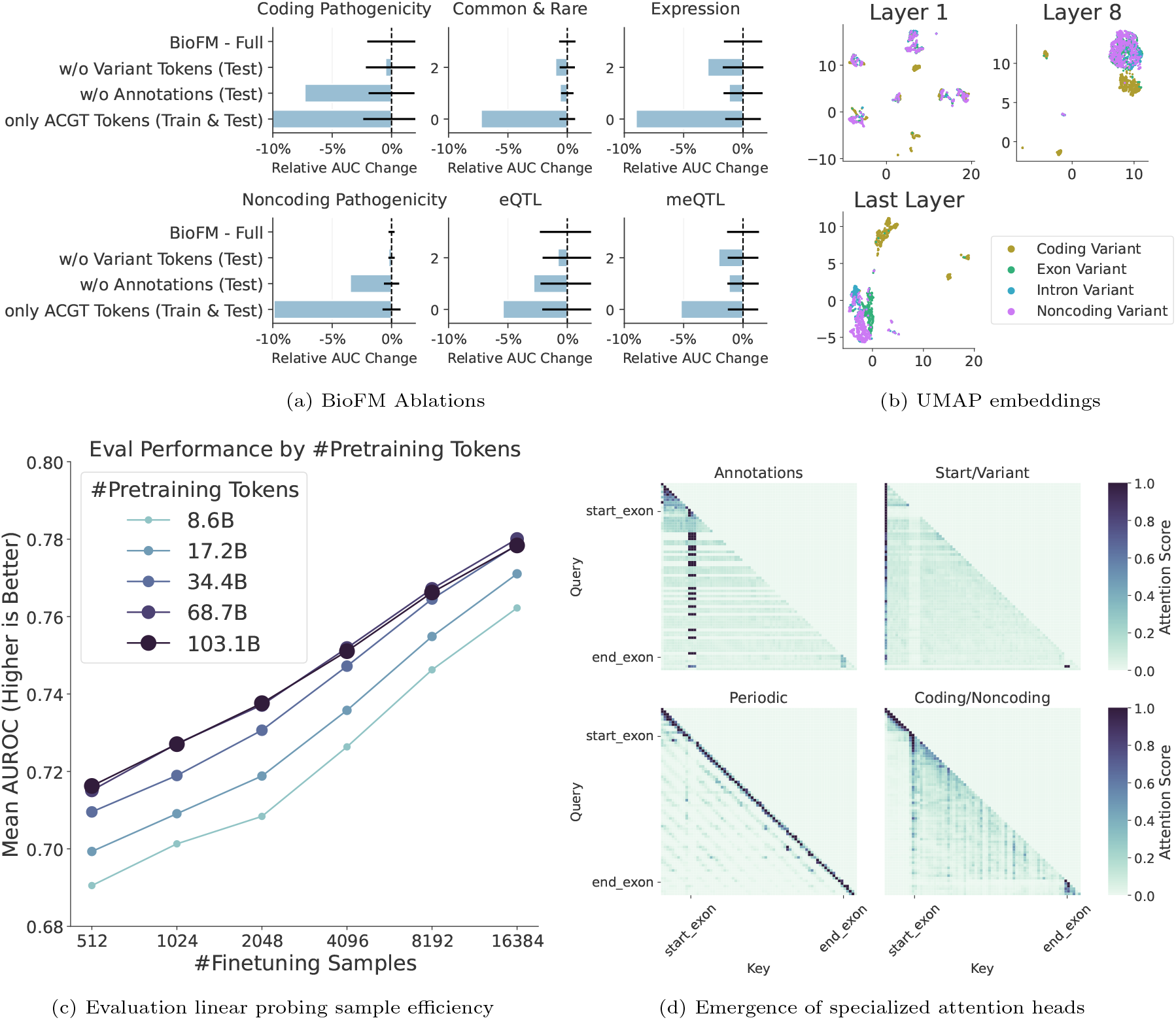
Detailed analysis shows that biologically-informed tokenization using BioToken in BioFM significantly enhances genomic modeling accuracy and computational efficiency. **a)** Ablation experiments clearly demonstrate that both annotation and variant-specific tokens in BioToken significantly contribute to improved genomic representation quality, with pronounced impacts on coding pathogenicity and splicing variant prediction. We removed annotations and variant tokens from test sequences and tested the original BioFM on them. Additionally, we trained a version of BioFM without annotation tokens and another version without annotation and variant tokens. Error bars are 2 standard deviations wide, centered on the mean. **b)** UMAP embeddings illustrate progressive differentiation of variant types through model layers, indicating BioFM effectively learns biologically meaningful distinctions such as coding versus noncoding variants. **c)** Evaluation of linear probing efficiency highlights BioFM’s sample efficiency and scalability, benefiting significantly from increased genomic training data while achieving superior downstream performance. **d)** Visualization of attention maps provides mechanistic insight into specialized attention heads, clearly demonstrating BioFM’s ability to selectively attend to biologically significant genomic annotations and regions, further validating the functional interpretability and efficacy of the BioToken strategy.

Next, we investigated the structure of variant representations learned by BioFM using UMAP embeddings derived from pathogenicity benchmark datasets (Figure 5b). This embedding analysis revealed progressive differentiation of variants in deeper model layers. While initial layers showed no biologically coherent separation, the final layer clearly distinguished coding from noncoding variants, with further sub-structuring between exonic and intronic variants. This aligns well with biological intuition, as coding variants primarily affect protein structure and function, whereas noncoding variants are predominantly involved in splicing and regulatory mechanisms.

We also evaluated BioFM’s ability to utilize large-scale pretraining well by analyzing its sample efficiency with linear probing on variant benchmark tasks. We expected the downstream task performance differences to be larger when the number of finetuning samples is low. In agreement with this hypothesis, we observed that the difference between undertrained (8.6B, 17.2B) and trained models (68.7B, 103.1B) is particularly pronounced when the number of finetuning samples is low (Figure 5c). The difference reduces when we increase number of fine-tuning samples for linear probing. This observation illustrates that there is a clear benefit to scaling training data size, with the law of diminishing returns eventually becoming apparent.

Lastly, we probed BioFM’s internal attention patterns to explore how biologically-informed annotations and variant tokens affect representation learning at a mechanistic level (Figure 5d). We hypothesized that specific attention heads may be specializing in processing annotation and variant tokens based on inherent sequence differences between coding and noncoding regions. This expectation aligns with the biological observation that the probability distribution of DNA k-mers differs between coding and noncoding regions–coding regions exhibit constrained sequence variation, while noncoding regions often contain repetitive elements. Attention mechanisms should thus help the model distinguish between these contexts by learning distinct probability distribution patterns.

To test this hypothesis, we selected a 64-base sequence that includes a short exon sequence in the middle, flanked by `⟨start_exon⟩ and ⟨end_exon⟩` tokens, with a variant token at the end. Visualization of attention maps for this sequence revealed distinct patterns among attention heads (Figure 5d). Some attention heads were dedicated to annotation tokens, while others focused on sequence start and variant tokens, potentially aiding in the recognition of contextual boundaries and mutation sites. Certain other attention heads exhibited periodic patterns, which may reflect the repetitive structure of noncoding regions, often associated with regulatory or structural genomic elements. Additionally, some attention heads segregated exonic and intronic regions, indicating a learned distinction between protein-coding and noncoding sequences. These findings indicate that the model recognizes biologically meaningful sequence features, offering insights into how BioToken benefits BioFM in encoding genomic information.

## Discussion and Conclusion

In this work, we introduced BioToken, a biologically-informed tokenization framework designed to improve the computational representation of genomic sequences and their variants. Using BioToken, we developed BioFM, a foundation model that achieves superior performance across a wide spectrum of downstream tasks. Unlike conventional genomic foundation models, BioFM explicitly encodes biological structures and variants into its token representations, enabling it to surpass existing GFMs of comparable size and even much larger models such as Evo2 7B across several tasks. Most significantly, BioFM matches or exceeds specialized models like Enformer and SpliceTransformer on their respective domain-specific benchmarks, bridging the longstanding gap between general-purpose genomic models and task-specific architectures. This makes BioFM the first GFM with a capability to model variant effects across the central dogma. We also showed that BioFM is competitive on region classification tasks and is able to generate promoters given a gene sequence.

The core innovation of BioFM stems from a fundamental shift in how genomic data is represented in machine learning models. Traditional approaches typically model DNA sequences as linear nucleotide strings devoid of biological context. BioToken corrects this limitation by encoding genomic variants (including SNVs, insertions, and deletions) and region annotations (such as exons, introns, transcripts, and coding regions) directly into specialized tokens. Our comprehensive ablation studies confirm that integrating these biologically meaningful annotations is the primary factor driving BioFM’s enhanced predictive performance.

A key strength of the BioToken framework is its modularity and extensibility. While our current implementation focuses on SNVs and basic genomic annotations, the framework can be readily extended to incorporate additional biological contexts. Future iterations could include tokens for chromatin 3D structure, methylation rates, histone modifications, or even cell- and tissue-type specific annotations. Another direction is to add information from single-cell foundational models [41–44] in special tokens similar to multimodal LLMs [45]. This extensibility ensures that BioToken can evolve alongside our growing understanding of genomic function and structure.

Another significant contribution of BioToken is its demonstration that encoding intrinsic biological biases directly into model inputs can dramatically improve computational efficiency without sacrificing performance. By providing the model with structured, biologically meaningful tokens, this method reduces the model’s reliance on implicit learning of genomic patterns, significantly boosting computational efficiency. BioFM achieved its results using only 333 billion training tokens compared to trillions of tokens used to train other models [3, 10]. Another relevant finding was that BioFM does not require the expensive task-specific layer-wise search for best representation, as the last-layer embedding from BioFM leads to superior or equivalent performance on downstream tasks compared to the intermediate layers. Together, these results make genome foundation modeling more accessible for the community.

Moreover, the principles underlying BioToken are not limited to DNA sequences. The same approach can be adapted for protein language models by augmenting traditional amino acid tokenizers with domain annotations and variant tokens. This could significantly enhance our ability to model protein structure-function relationships and predict the functional consequences of amino acid substitutions. The potential applications extend to multi-modal models that integrate DNA, RNA, and protein data within a unified representation framework.

While our results are promising, several limitations merit consideration. First, the current implementation of BioToken and BioFM relies on high-quality annotations and variant data, which may not be equally available across all organisms. Application to non-model organisms or species with limited genomic resources would require adaptation and potentially alternative approaches to annotation. Second, the scaling properties of our model with increased variant data remains open for future works. While we trained on data from 50 individuals from the 1000 Genomes project, extending to the full scale of this dataset (nearly 3200 whole genomes) or biobank-scale data [46] could provide significant gains with a proportional increase in the model size. Third, we recognize that annotation tokens may introduce systematic biases during evaluation. For example, eQTLs and sQTLs are, on average, closer to transcript or exon boundaries than negative controls. To address this, we conducted experiments with distance-matched negative controls and found that performance decreased across all models, yet BioFM consistently maintained the strongest results (Figure 2g). Furthermore, several of our tasks incorporate such disntance-matched negative controls by design. The overarching premise of large-scale foundational pretraining, as outlined in Evo2 [10], is that models should eventually internalize the structure of genes, exons, and coding regions. From this perspective, providing accurate annotation tokens offers a more efficient alternative to forcing the model to learn these features indirectly. At the same time, our findings suggest that incorporating available genomic annotations into sequence-based models may facilitate more effective learning and prioritization of relevant features, although the extent of this benefit warrants further investigation.

In conclusion, BioToken and BioFM represent a meaningful advance in genomic modeling, providing a robust and extendable framework for modeling the complex relationships between genetic variation and molecular phenotypes. This work establishes a new paradigm for genomic foundation models that prioritizes biological context and computational efficiency, offering a promising path forward for the field of computational genomics and its applications in precision medicine.

## Code and Model Availability

- BioFM model is available on Hugging Face: https://huggingface.co/m42-health/BioFM-265M
- Python package for inference and embedding extraction is available on GitHub: https://github.com/m42-health/biofm-eval/
- Variant benchmark is available on Hugging Face: https://huggingface.co/datasets/m42-health/variant-benchmark
- The training code is adapted from Meta’s llama-cookbook: https://github.com/meta-llama/llama-cookbook

## Data Availability

- The BioFM pre-training sequences were obtained from publicly available resources. The 1000 Genome Project sequences were obtained form https://ftp.1000genomes.ebi.ac.uk/vol1/ftp/data_collections/1000G_2504 high coverage, and the human reference genome from https://www.ncbi.nlm.nih.gov/datasets/genome/GCF_000001405.26/.
- Gene annotations were obtained form GENCODE, https://www.gencodegenes.org/human/release_38.html.
- The process of creating variant benchmark and the processed data is made available on Hugging Face: https://huggingface.co/datasets/m42-health/variant-benchmark which encompasses tasks like coding and non-coding pathogenicity assessment, expression effect prediction, alternative splicing, DNA methylation, ancestry classification, and common vs. rare variant prediction.
- Genomic long range range benchmark is collected from https://huggingface.co/datasets/InstaDeepAI/genomics-long-range-benchmark.
- GUE benchmarks are prepared using the official GitHub repo from authors: https://github.com/ML-Bioinfo-CEITEC/genomic_benchmarks.
- The data used for regulatory sequence analysis is collected from https://epd.expasy.org/epd.

## Appendix A Methods

### A.1 Data

#### A.1.1 Pretraining Dataset

We utilized whole genome sequencing (WGS) data from the 1000 Genomes Project [14] as the pretraining dataset. From the total pool of 3,202 available samples, we selected a subset of 50 individuals. This subset was balanced across the five superpopulations defined in the 1000 Genomes dataset, with 10 individuals randomly chosen from each superpopulation. The list of individuals is in table Table A1. We used Phase 3 WGS data in HG38 assembly. Variants are phased as described in [14].

During training, we generated input sequences by sampling genomic regions of 6,144 nucleotides in length. To increase data diversity and ensure overlap between sequences, we introduced a sliding window approach with an overlap of 1,000 nucleotides between consecutive regions. Additionally, for each sequence, we applied a random start shift uniformly sampled between 0 and 1,000 nucleotides to further vary the input segments. As data is phased, each time we randomly selected one of two haplotypes.

We excluded any 6,144-nucleotide region in which more than 50% of the bases were unknown (i.e., represented by ‘N’) in the human reference genome, to ensure data quality. For the remaining sequences, we introduced sample-specific genetic variation. Specifically, for each selected region, we randomly chose one of the 50 individuals and applied all of their corresponding single nucleotide variants (SNVs) and insertions or deletions (indels) that mapped to that region. This process was expected to yield a mutated sequence reflecting real human genetic diversity. The training process covers each region for each individual exactly twice. The total number of regions is 542,688, each of them is seen by the model 100 times, yielding total number of tokens equal to 333B.

To standardize input length for the model, each mutated sequence was truncated or padded to exactly 6,144 tokens. We further applied reverse complement augmentation with a probability of 50% for each region. When reverse complementing, transcript, exon, and coding sequence boundary annotations were flipped only if the original strand corresponded to the forward strand; otherwise, they were left unchanged to preserve correct strand-specific annotation. Pretraining was performed using data from autosomes (chromosomes 1 to 22) and the X chromosome.

**Table A1:** 50 samples used for pretraining BioFM.

| AFR | EUR | SAS | EAS | AMR |
| --- | --- | --- | --- | --- |
| NA19835 | NA20506 | NA20902 | HG02040 | HG01281 |
| HG03571 | HG00251 | HG04001 | HG01808 | HG02312 |
| NA19258 | HG01707 | HG03899 | NA18637 | HG00639 |
| HG03055 | HG00365 | HG03905 | HG01858 | HG01374 |
| NA18930 | NA12872 | HG02662 | HG01878 | HG01950 |
| NA19320 | NA12812 | HG03703 | HG00706 | HG01974 |
| NA19035 | HG00159 | HG04176 | HG00472 | HG01178 |
| HG02307 | HG00272 | HG02649 | HG01597 | HG02285 |
| HG03086 | NA12275 | HG03885 | NA18608 | HG02292 |
| NA19379 | HG01771 | HG03778 | HG00694 | HG01200 |

##### A.1.2 Evaluation Datasets

###### Noncoding Pathogenicity

This task involves classifying genetic variants located in noncoding regions of the human genome as either pathogenic (disease-causing) or benign (non-disease-causing). We use the BEND benchmark dataset [22], derived from ClinVar [47] and includes single nucleotide polymorphisms (SNPs) annotated with binary pathogenicity labels. Specifically, the dataset comprises 295,923 SNPs, of which 21,524 are labeled as pathogenic and 274,399 as benign, resulting in a significant class imbalance ( 5% pathogenic). All variants are mapped to the GRCh38 reference genome.

To construct BEND, the authors first filtered ClinVar (variant summary file dated 2023-07-02) for single nucleotide variants with at least one-star review status. To isolate noncoding SNPs, they removed variants annotated as coding in ClinVar, and further excluded SNPs located in coding sequences (CDS), start codons, and stop codons based on GENCODE v43 annotations [48].

Variants from the mitochondrial genome were also omitted to maintain consistency with prior literature baselines. Pathogenicity annotations were binarized by combining the “Likely pathogenic” and “Pathogenic” categories into a single pathogenic class, and “Likely benign” and “Benign” into a single benign class, following [48]. Additionally, the authors applied Ensembl VEP [49] to categorize variants by consequences. The original task from BEND benchmark is designed as a zero-shot classification task, but in this work, we applied it in a supervised classification setting. Further details on the evaluation procedure are provided in Appendix A.3

The current state-of-the-art performance on this benchmark is achieved by CADD v1.7, which reports an AUROC of 0.992 [50]. It is important to note that CADD is an aggregate scoring method that integrates multiple predictive features, some of which may incorporate signals from models trained on ClinVar data.

###### Coding Pathogenicity

The coding pathogenicity prediction task focuses on classifying genetic variants located within coding regions of the genome as either pathogenic (disease-causing) or benign (non-disease-causing). Coding regions are responsible for protein synthesis and are therefore central to understanding disease mechanisms, making this task a critical component of medical genomics and precision medicine.

For this task, we use a dataset derived from the AlphaMissense [26], which filters and curates missense variants from ClinVar (release date: 2022-09-10). Variants were mapped to the canonical transcript, and those labeled as “Benign” or “Likely benign” in ClinVar were treated as benign, while those labeled as “Pathogenic” or “Likely pathogenic” were treated as pathogenic. Variants containing the term “splice” in their annotation were removed (29 variants), as the focus of this task is on missense mutations rather than splicing effects. Following this processing, a total of 100,796 missense variants with at least one review star were considered.

The final test dataset consists of 82,872 variants from 7,951 proteins, after removing those used for validation. Of these, 30,884 are pathogenic and 51,988 are benign. This dataset reflects real-world class imbalance but remains significantly more balanced than typical noncoding variant datasets, offering a robust benchmark for model evaluation. Moreover, coding variants are generally better annotated and more frequently subjected to functional validation, which contributes to higher data quality in this task.

Importantly, variants in coding regions frequently lead to missense mutations that directly alter protein structure or function. While traditional tools such as CADD [50] have been widely used for this problem, novel GFM approaches, such as GPN-MSA, are beginning to surpass conventional methods in predictive performance.

In this study, we adopt the test set proposed in the AlphaMissense paper for both model training and evaluation to assess coding pathogenicity performance. AlphaMissense has AUROC of 0.940. Further details on the evaluation procedure are provided in Appendix A.3.

###### eQTL Prediction

This task involves predicting whether a genetic variant influences gene expression. The ability to predict expression-altering variants has far-reaching implications for understanding the genetic basis of traits and diseases. Accurate predictions can guide experimental validation, prioritize variants for functional studies, and inform the development of gene therapies.

The dataset used for this task was adapted from the Borzoi study [23] and consists of tissue-specific VCF files for 41 human tissues (filtered from the original 49 due to too few data samples in 8 tissues), publicly available at https://console.cloud.google.com/storage/browser/borzoi-paper/qtl/eqtl. The balanced dataset comprises a total of 97,922 variants, consisting of half positive eQTLs and an equal number of matched negative controls. The positive set includes variants with a posterior inclusion probability greater than 0.99 following SuSiE fine-mapping [51]. To control for genomic proximity as a confounding factor, negative controls were specifically chosen to match the positive eQTLs in their distance to the transcription start site (TSS), following the procedure detailed in the original study. For the linear probing evaluation on Variant Benchmark, a separate logistic regression model was trained for each fold within each tissue. Performance was evaluated using an 11-fold cross-validation scheme. AUROC was calculated for each fold and subsequently averaged to yield a mean AUROC for each tissue. A final aggregate AUROC and standard deviation were then computed by taking a weighted average of the per-tissue results.

###### Expression Prediction

The dataset for this task was derived from DeepSea [24]. Variants with positive labels were obtained from the GRASP eQTL database [36], lifted over from hg37 to hg38 using PyLiftOver (https://pypi.org/project/pyliftover/), and filtered to exclude conversion errors.

To construct a balanced dataset, each positive variant was paired with a randomly selected negative control from the original dataset, resulting in 156,162 variants in total.

To further investigate the role of TSS distance to variants, we created two additional datasets corresponding to *<*360 bp and *<*31 kbp negative control sets, as outlined in Figure 2g. For the *<*360 bp version, negative controls were restricted to variants within 360 bp of a positive sample, following the evaluation procedure in DeepSea. This subset included 57,457 negative controls, compared to 78,081 negatives in the default balanced dataset. For the *<*31 kbp version, we combined the *<*360 bp set with additional negatives within 710 bp, 6.3 kbp, and 31 kbp of positive variants, using the original DeepSea controls.

These additional configurations for datasets were designed to test whether BioFM performance relies on variant-TSS distance information encoded by annotation tokens. This process enables a clearer assessment of whether model performance reflects genuine learning of regulatory effects rather than dependence on distance-based signals.

###### sQTL Prediction

The splicing effect prediction task aims to determine whether a genetic variant influences alternative splicing patterns. Evaluation splicing effects is critical for understanding the mechanisms by which variants contribute to diseases, particularly for disorders caused by aberrant RNA processing. Existing methods primarily focus on predicting splicing sites, rather than the direct effects of specific variants on splicing, leaving a gap in accurate variant-level predictions.

The dataset for this task was obtained from the Borzoi study [23], comprising tissue-specific VCF files with positive sQTLs and matched negative controls (https://console.cloud.google.com/storage/browser/borzoi-paper/qtl/sqtl ), following a similar procedure as for the eQTL dataset. In total, 43,028 variants were collected. Initial evaluations revealed that several tissue-specific folds contained very few variants, in some cases lacking either positive or negative samples. To address this limitation and ensure robust evaluation, tissues were grouped into broader categories (brain, hematopoietic, digestive, urogenital, adipose, endocrine, and others). Within each cluster, duplicate variants across tissues were removed to avoid redundancy (e.g., overlapping positive sets from multiple brain tissues). Variant counts for each tissue cluster are reported in Table A2. Evaluation was performed by computing AUROC for each fold within a cluster, followed by a weighted average across folds to obtain the final AUROC and standard deviation.

**Table A2:**
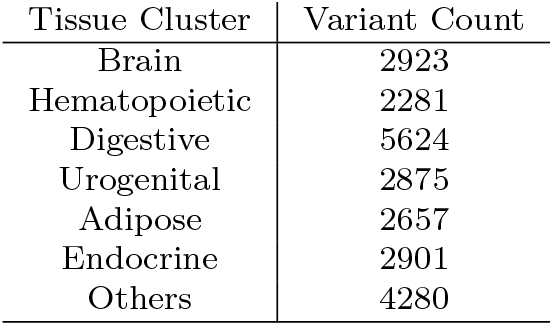
sQTL statistics.

###### Ancestry Prediction

The ancestry prediction task aims to classify individuals into one of five superpopulations based on genomic sequence data. To construct the dataset, we started with the human reference genome (HG38) and generated consensus sequences for each individual in the 1000 Genomes Project [14] by applying their corresponding mutations. For each individual, 11 genomic regions were selected, with each region comprising 32,000 nucleotides. These regions are sampled from alternate chromosomes (e.g., chromosomes 1, 3, 5, etc.), and their positions are fixed across all individuals in the dataset. Variability arises exclusively from the individual-specific mutations applied to these regions.

On average, the sequences differ from the reference at approximately 0.5% of positions, corresponding to around 33 variants per sequence. These variants include single nucleotide polymorphisms, insertions, and deletions. To ensure data quality, the selected regions are constrained such that all individuals in the 1000 Genomes dataset are fully represented in each region, i.e., there are no ambiguous or unknown bases, and the regions exhibit sufficient genetic variation across samples.

In the final dataset, the 11 regions are treated as independent samples, and each is assigned a superpopulation label corresponding to the individual from whom it was derived. This design results in a dataset that is 11 times larger than the number of individuals in the original 1000 Genomes dataset.

For model training, each 32,000-nucleotide sequence is divided into smaller chunks to accommodate model-specific context length limitations (e.g., 6,144 for our BioFM, 8,192 for Evo2-1B-Base, and 12,288 for NT). Embeddings are extracted for each chunk, and a mean pooling operation is applied to obtain a single representation for the entire sequence. These aggregated embeddings are then used as inputs to a linear classifier to predict ancestry.

###### Common vs Rare Variants Classification

This task assesses whether model embeddings capture biologically conserved genomic contexts typical of authentic common variants. We randomly sampled 100,000 common variants (Minor Allele Frequency (MAF) *>* 0.05) from GnomAD [15] and, for each common variant, generated a synthetic negative control by randomly substituting a nucleotide within a ±16-nucleotide local sequence window. The resulting balanced dataset consists of exactly 200,000 single-nucleotide variants (SNVs), equally divided between genuine common variants and synthetic controls.

Generation of synthetic negative controls in a close proximity to common variants ensures that annotation tokens do not leak the information into the testing set. Moreover, during evaluation stage, both common variants and negative control variants are encoded with a variant token, therefore this task tests models ability to generate different embeddings for common and rare variants.

###### meQTL Prediction

This task involves predicting if a genetic variant influences nearby methylation rates. We downloaded meQTL dataset from GRASP [36] which contains 52,418 datapoints. We filtered out duplicates by rsid, lifted variant coordinates to hg38 and obtained 25,496 variants (meQTLs) total. GRASP meQTL dataset itself is a compilation of meQTLs found by many studies, some of them are verified experimentally, others are fine-mapped. As this dataset contains only positive samples, to create negative controls, we randomly sampled 100,000 common variants (Minor Allele Frequency (MAF) *>* 0.05) from GnomAD [15], grouped them by chromosome and GC-content percentage with an precision of 1%, and for each meQTL variant we selected a random negative control from the corresponding group with the same chromosome and GC-content percentage. This procedure ensures that positive samples and negative controls are not easily distinguishable by GC-content around variant.

#### A.2 Modelling

**Table A3:**
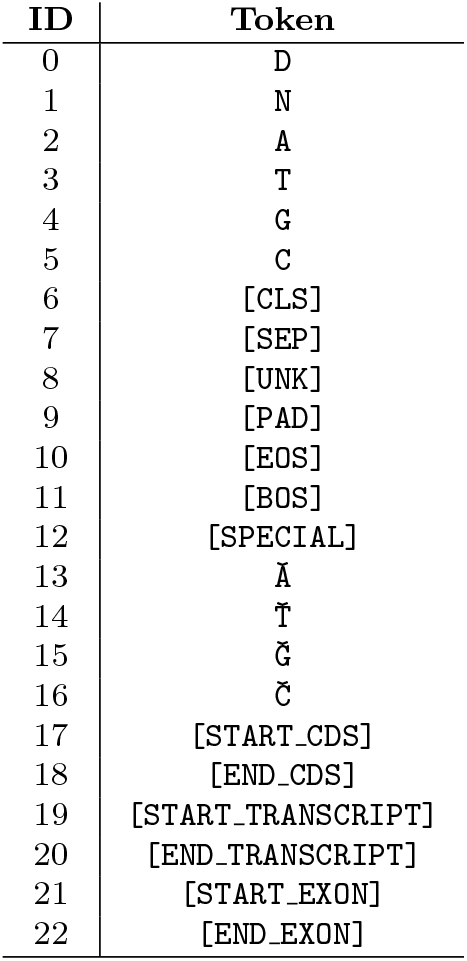
BioToken Annotation Variant Vocabulary.

##### A.2.1 BioToken

We experiment with three variants of BioToken to explore varying levels of genomic complexity. Each tokenizer incorporates unique aspects of genomic data, enhancing model understanding and performance. Figure 1a provides an overview of nucleotide-level points of interest and their integration into tokenization strategies. A brief description of the tokenizers is provided below:

###### BioToken-Base

This baseline tokenizer maps each nucleotide (A, T, G, C) to a distinct token. It is commonly used in genomic foundation models [4] and offers a simple yet effective representation that preserves base-level positional information and facilitates preprocessing. However, this character-level encoding treats all genomic positions equivalently, ignoring biological context or inductive biases. Unlike natural languages, DNA sequences convey meaning not through syntactic rules, but through biochemical mechanisms, spatial chromatin structure, and regulatory interactions. As a result, modeling the genome purely as linguistic text overlooks its intrinsic organization and biological regularities.

###### BioToken-Variant

To capture individual-specific genetic variation, this tokenizer extends the BioToken-Base by incorporating variant information relative to the reference genome. For each individual from the Genome1000 dataset, variants within the sampled genomic regions are explicitly identified and encoded. Special symbols—Ă, Ť, Ğ, Č—are used to represent single nucleotide variants and insertions. This design enhances the model’s ability to detect and learn from regions of genomic diversity.

###### BioToken–Annotation-Variant

Building upon the BioToken-Variant, this approach incorporates additional functional annotations crucial for understanding genomic patterns. Annotations include start and end of coding sequences, transcription regions, and exon regions providing context about regions critical to gene expression. These annotations enable the model to better capture biologically significant patterns. We use the GENCODE annotations version 38 [52]. We retain only CDS, transcript, and exon annotations, and remove exact duplicates based on annotation type and start/end coordinates. If two annotations of the same type share the same start position, both are included in sequence (e.g., two consecutive start exon tokens). It is important for the model to know the extent of alternative splicing of genes. The vocabulary for the annotation variant tokenizer is provided in Table A3.

##### A.2.2 BioFM

We utilize a decoder-only transformer architecture derived from Mistral [53] as the foundation for our model. The architecture consists of 23 transformer layers with a total of 265M parameters. While we retained the grouped-query attention mechanism from the original Mistral architecture to improve inference efficiency, we disabled the sliding-window attention feature to allow the model to access the full input context without local attention constraints. To accommodate longer genomic sequences, we modified the Rotary Position Embeddings (RoPE) by increasing the *θ* parameter from 10^4^ to 10^5^, following established approaches for extending context length in transformer models [33]. This modification enables the model to better handle positional information across extended DNA sequences that are typical in genomic applications. All remaining hyperparameters, including learning rate, batch size, and optimization settings, are detailed in Table A4.

#### A.3 Evaluation

##### Variant Benchmark

For each task in Variant Benchmark, except ancestry prediction, we divide variants into 11 folds. Each fold has 21 chromosomes as training set and 2 chromosomes as testing set, except fold 0, which has chromosomes 21, 22, and X as testing set. For encoder models, we put variant in the center of context window, construct embeddings of reference sequence and mutated sequence and concatenate them. For decoder-only models, we construct forward mutated sequence which ends with variant, and reverse complement mutated sequence which also ends with variant. Both forward and reverse sequences have length of exactly half of evaluation context size. After that, we average forward and reverse reference sequences, and forward and reverse mutated sequences, and concatenate the resulting two vectors. This approach ensures that each model has access to the exact same amount of context information and also works around the fact that decoder-only models are causal.

After obtaining embeddings for each variant, we concatenate them into a feature matrix for a linear regression. We call it linear probing. We decided against full or parameter-efficient finetuning to mitigate the overfitting effects and effect of many additional degrees of freedom coming from having many hyperparameter combinations to select from. Linear probing of model embeddings also allows us to isolate the quality of obtained representations. We used default settings in a sklearn package, but with number of steps for lbgfs method set to 5000. We report AUROC for each test set in each fold, and then obtain average AUROC for Figure 2a or median AUROC for Figure 1d and **??**. It allows us to get more robust performance estimates for each model.

**Table A4:** Hyperparameters for BioFM 265M.

| Parameter | Value |
| --- | --- |
| optimizer | AdamW |
| max_learning_rate | $7.2e - 4$ |
| min_learning_rate | $3.2e - 5$ |
| weight_decay | 0.1 |
| optimizer_momentum | $\beta_1, \beta_2 = 0.9, 0.999$ |
| batch_size | 64 |
| context_size | 6144 |
| lr schedule | cosine with warmup |
| epochs | 2 |
| initializer_range | 0.03 |
| hidden_size | 640 |
| n_layers | 23 |
| n_heads | 32 |
| n_gqa | 8 |

For comparison with Enformer, we use a PyTorch Enformer implementation from EleutherAI github.com/lucidrains/enformer-pytorch. Enformer first compresses each sequence into blocks using convolutions. Following original evaluation approach in [27], we extract three hidden states for blocks centered about variant of interest, average them into one hidden state over sequence dimension and use it for the linear probing. Additionally, we ran an experiment where we used only block with the variant, but results did not change. We evaluated Enformer on expression prediction task from Variant Benchmark using context sizes of 12K, 24K, 48K, and 96K. Performance of Enformer saturated for 48K and 96K nucleotides, therefore we did not evaluate it on 192K context.

For comparison with Splice Transformer, we use an official implementation from authors at github. com/ShenLab-Genomics/SpliceTransformer. We always use the maximum available context size of 8K nucleotides, centered about variant of interest, extract embeddings for this variant only, and use linear probing on top.

For comparison with Evo2-1B and Evo2-7B-Base models, we use embeddings from layer 19 and layer 24, respectively. We selected these layer indices, because embeddings from last layer performed significantly worse and in [10] these layer embeddings were generally among the best for coding pathogenicity prediction in BRCA1 and BRCA2 tasks. We also use mean pooling for all results shown in Figure 2a. It consistently shows better performance compared to last token pooling. We also evaluated context sizes (sequence length) for both last token pooling and mean pooling from 1024 to 8192, and on average, mean pooling with sequence length of 1024 works better. Therefore, to create a maximally favourable conditions for Evo2 set of models, we use specific layer embeddings, mean pooling, and context size of 1024.

##### GUE Evaluation

We use the existing train-validation-test splits on the original GUE dataset for finetuning purposes. For each model, we ran a sweep through 4 values of learning rate and 2 values of maximum epochs. The validation dataset is used for model selection, and the test scores corresponding to model with the best validation score is reported. As stated in Vishniakov et al., we find that max-pooling performs better than cls/last token pooling during our initial experiments [18]. Our finetuning configuration is provided in Table A6.

##### Dart Evaluation

We selected two tasks from Dart-Eval benchmark [31], namely African caQTL prediction and Yoruba dsQTL prediction. As original datasets are in HG37 assembly, we lifted them to HG38 assembly and filtered out variants with reference allele mismatch. After liftover, number of variants in African caQTL dataset is 79,026, and number of variants in Yoruba dsQTL dataset is 27,373. Prevalence of caQTLs in African dataset is 0.086, and prevalence of Yoruba dsQTLs is 0.02.

**Table A5:**
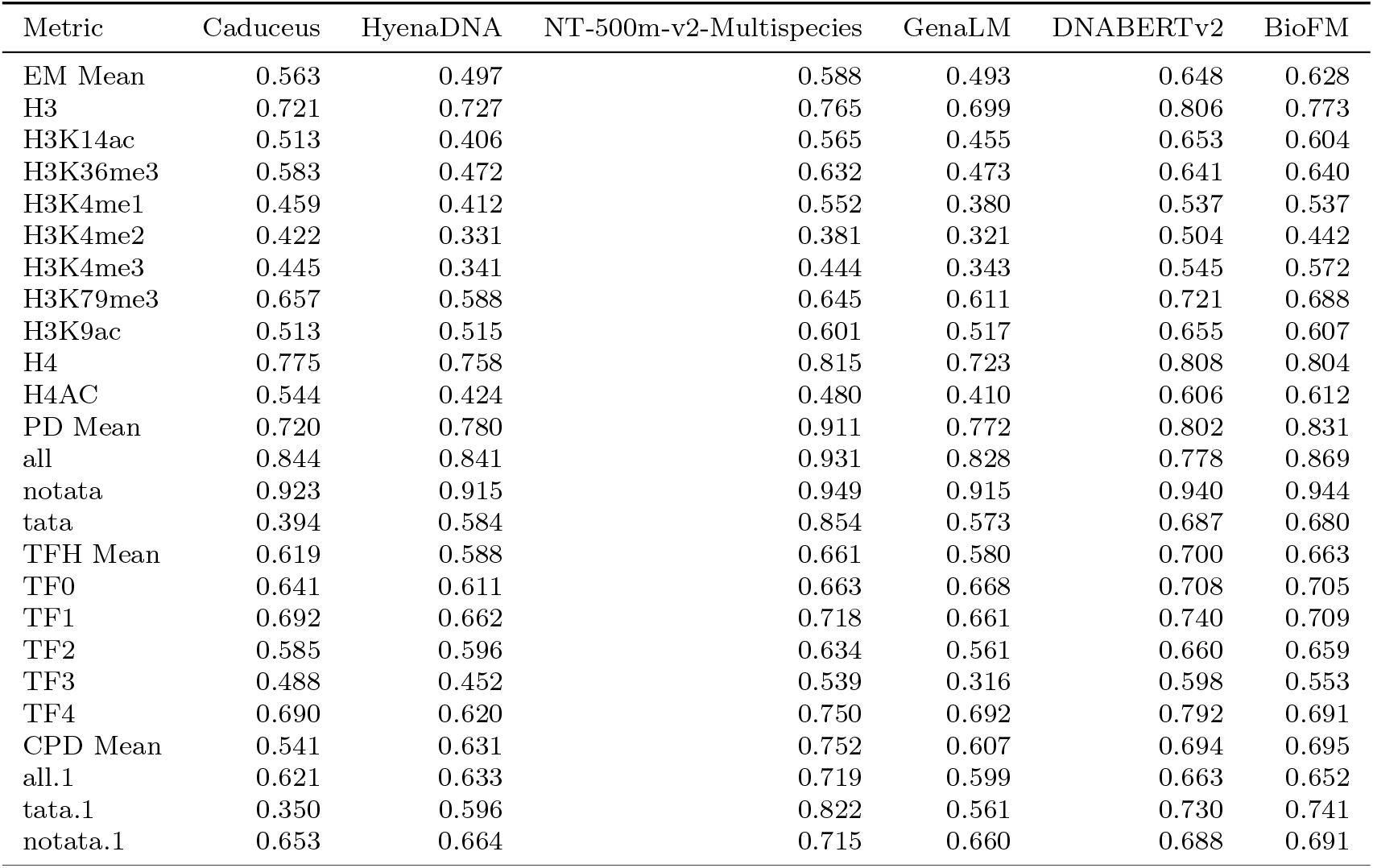
Full GUE Results.

**Table A6:** Configuration for GUE finetuning.

| config | value |
| --- | --- |
| optimizer | AdamW |
| learning_rate | 3e-5, 5e-5, 7e-5, 1e-4 |
| weight_decay | 0.01 |
| warmup_steps | 40 |
| optimizer_momentum | $\beta_1, \beta_2 = 0.9, 0.999$ |
| batch_size | 32 |
| lr schedule | cosine |
| epochs | 10 / 20 |

We used the same evaluation procedure as for our own Variant Benchmark, splitting data into 11 folds by chromosome and performing linear probing on top of every model embeddings afterwards. We used last layer embeddings and maximum context size up to 12K for every model, except Evo2 models, where we used context size of 1K and layers 19 and 24 for Evo2-1B-Base and Evo2-7B-Base, respectively. We chose to report AUROC to be consistent with choice of other benchmarks metrics.

##### Trait Gym Evaluation

Trait Gym has two variant-related tasks, called Complex traits prediction and Mendelian traits prediction. Both tasks include binary classification of variants which can influence a trait. The detailed variant selection procedure is described in [30]. Complex traits dataset has 10K variants, and Mendelian traits dataset has 3380 variants. We are using the same evaluation protocol, as in [30]. More specifically, for complex traits dataset, we split data into 22 folds, one fold per chromosome. We use linear probing with Elastic-Net regularization using GridCV search for the regularization parameter *C* from scikit-learn library. The *C* parameter grid is from 10^−8^ to 10^10^, similar to [30]. For the selection of the best value of *C*, we split training part of the dataset into 5 folds with train and validation parts and select *C* based on validation part of the dataset. Final reported metric is a weighted average AUROC, calculated on test splits of the dataset. We used last layer embeddings and maximum context size up to 12K for every model, except Evo2 models, where we used context size of 1K and layers 19 and 24 for Evo2-1B-Base and Evo2-7B-Base, respectively.

##### Promoter Generation

We use a high-quality experimental dataset EPDNew [40] for the Supervised Fine-Tuning (SFT). It contains 20,172 promoters for 9K genes. We split this dataset into 90% train genes and 10% test genes. After constructing gene sequences and accounting for reverse strands, we annotate each gene sequence with relevant metadata corresponding to the promoter dataset. Promoters are extracted using a − 249,+50 bp window around the transcription start site, following the approach used in the GUE dataset [54]. Additionally, we collect gene-specific information, including gene symbol, start, and end positions, to provide better contextual representation of the generated promoters. We fine-tune our pre-trained model for promoter generation using the SFTTrainer from the TRL library [55]. Our training data follows the structured response template: *<*gene sequence*>* P *<*promoter sequence*>*, which serves as inputs for the SFTTrainer.

##### Ablation, Scaling, Representation Analysis, and Attention Specialization

We conducted detailed analyses to better understand BioFM’s internal representations and the influence of training scale on downstream task performance.

For representation analysis, we constructed two-component UMAP embeddings of variant representations from the smaller 134M-parameter BioFM model, which contains 15 layers. Sequences used for embedding were fixed at 1024 nucleotides in length for both forward and reverse strands, with each variant consistently positioned at the sequence end. We averaged embeddings from forward and reverse strands to obtain the final variant representation. UMAP embeddings were computed using the Python umap package, with parameters set to minimal distance 0.05 and number of neighbors 10. To assess biological distinctions captured by these embeddings, we subsampled 500 variants for each Ensembl VEP consequence type from pathogenicity datasets within the Variant Benchmark.

To evaluate how downstream linear probing performance scales with the number of training tokens and finetuning samples, we subsampled only the training sets from each fold across seven tasks in the Variant Benchmark. The test sets remained constant throughout these evaluations. We considered five model checkpoints trained on varying amounts of genomic data, ranging from 8.6*B* tokens (equivalent to approximately 2.5 genomes) to 103.1*B* tokens (approximately 35 genomes). For each subsampled training dataset, we computed the average AUROC across the full test sets of each task, thereby systematically quantifying the relationship between pretraining scale and downstream prediction accuracy.

Finally, to investigate attention specialization, we analyzed attention maps using a fixed 64nucleotide sequence containing a small exon and coding sequence (CDS) at its center. We examined attention heads specifically selected for their hypothesized biological relevance: annotation and start/variant token heads from the initial layer (layer 0), and periodic and coding/noncoding heads from the final layer. This analysis aimed to provide mechanistic insights into how BioFM learns biologically meaningful patterns at the nucleotide level.

**Table A7:**
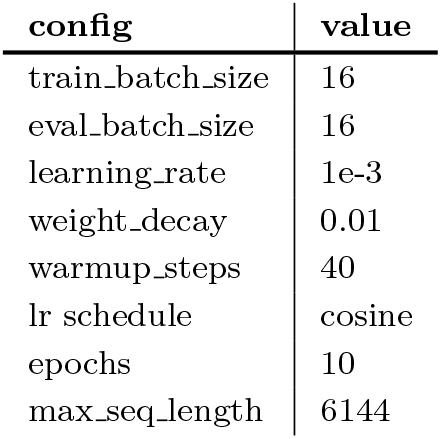
Configuration for SFT step.

| config | value |
| --- | --- |
| train_batch_size | 16 |
| eval_batch_size | 16 |
| learning_rate | 1e-3 |
| weight_decay | 0.01 |
| warmup_steps | 40 |
| lr schedule | cosine |
| epochs | 10 |
| max_seq_length | 6144 |

**Fig. A1:**
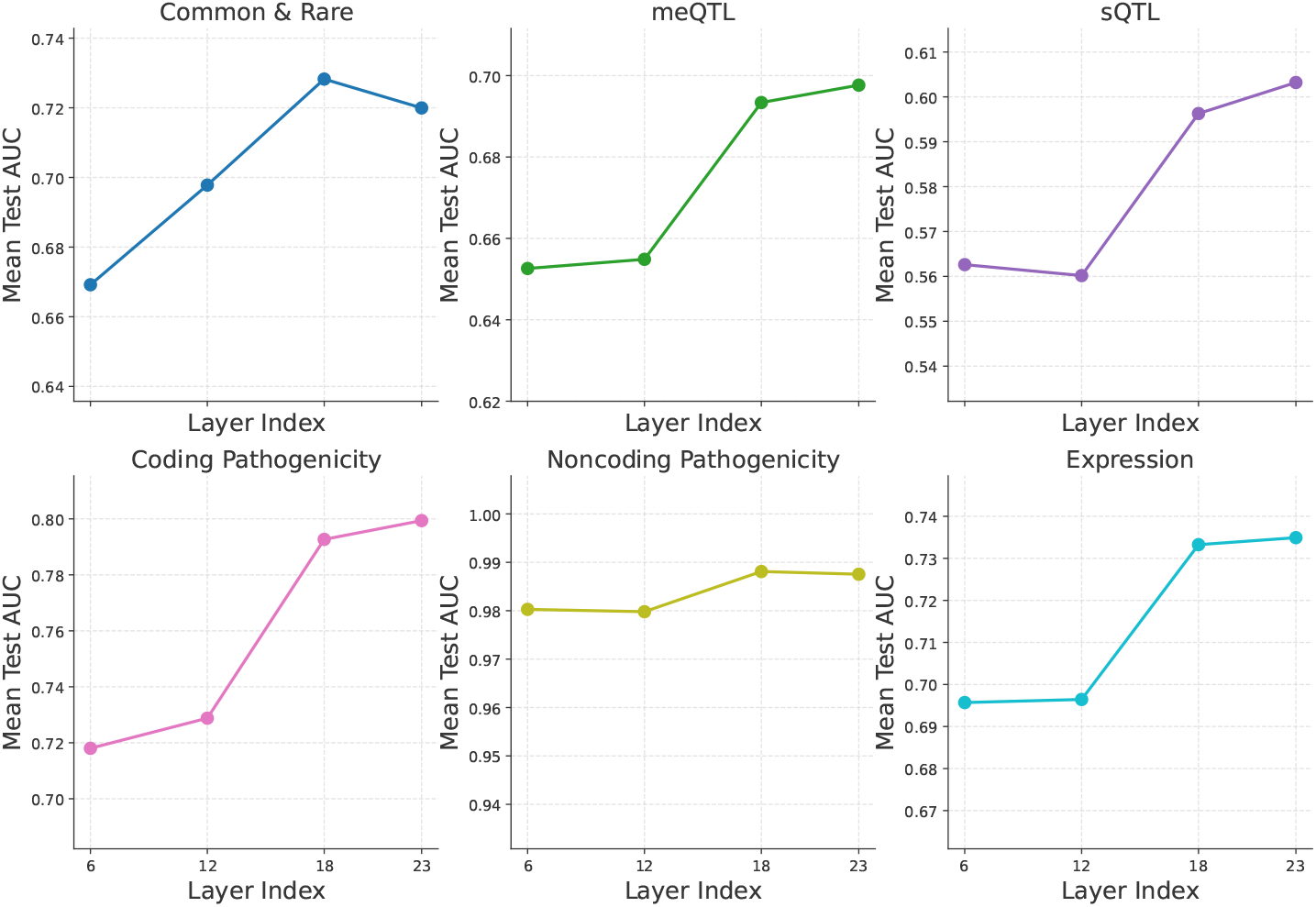
Different layer embeddings performance of BioFM 265M.

**Fig. A2:**
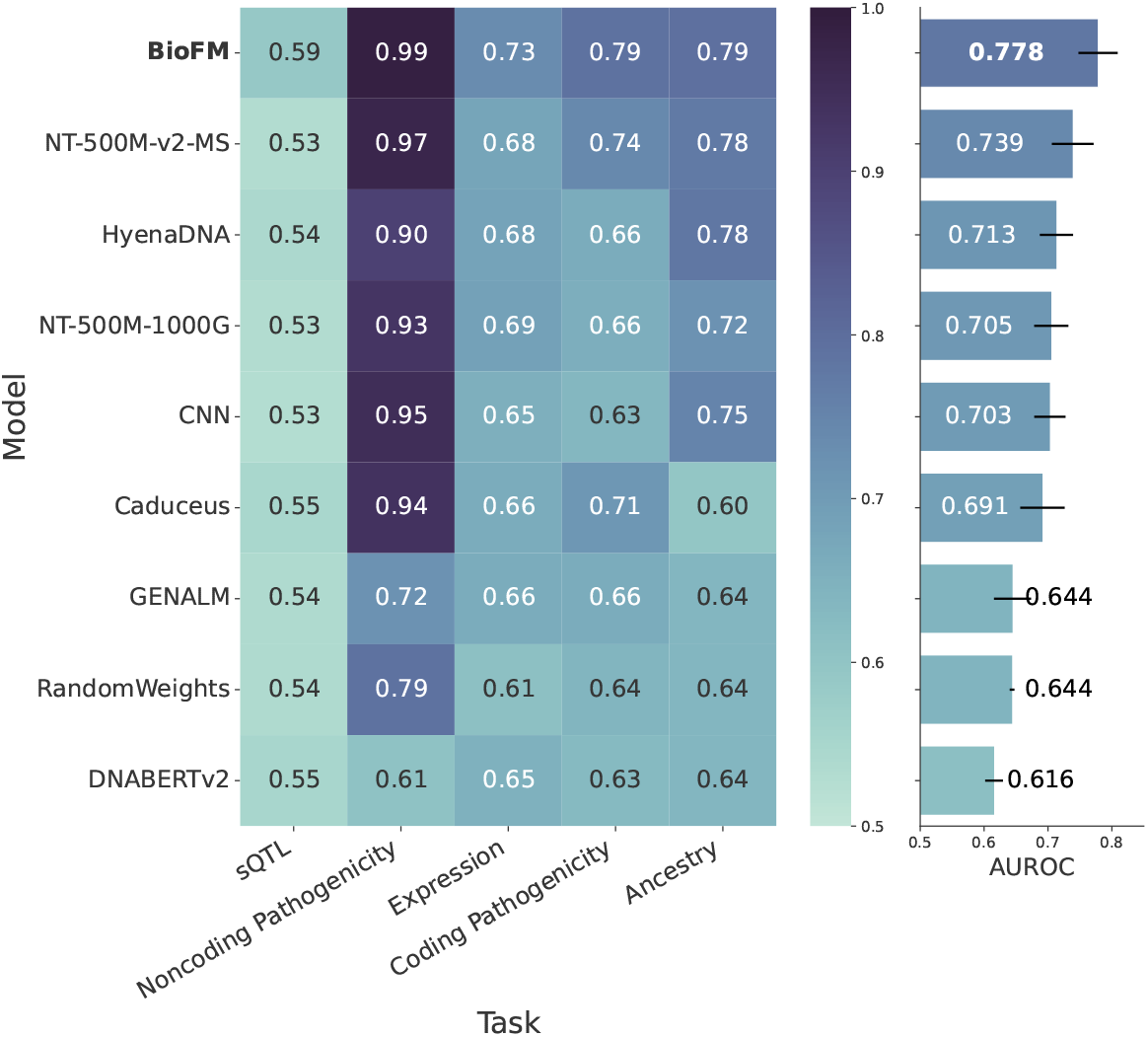
Variant Benchmark linear probing GFM comparison using context length of 1K. RandomWeights model correspond to the untrained version of BioFM 134M, which essentially produces random embeddings.

#### A.4 Pseudocode for BioFM Inference and Embedding Extraction

This pseudocode illustrates how BioFM-Eval Python library processes VCF files to generate biologically informed embeddings using the BioFM model. It includes variant annotation, context-aware sequence construction, tokenization, and final embedding extraction.

##### Algorithm 1

Variant Embedding Extraction Pipeline in BioFM-Eval

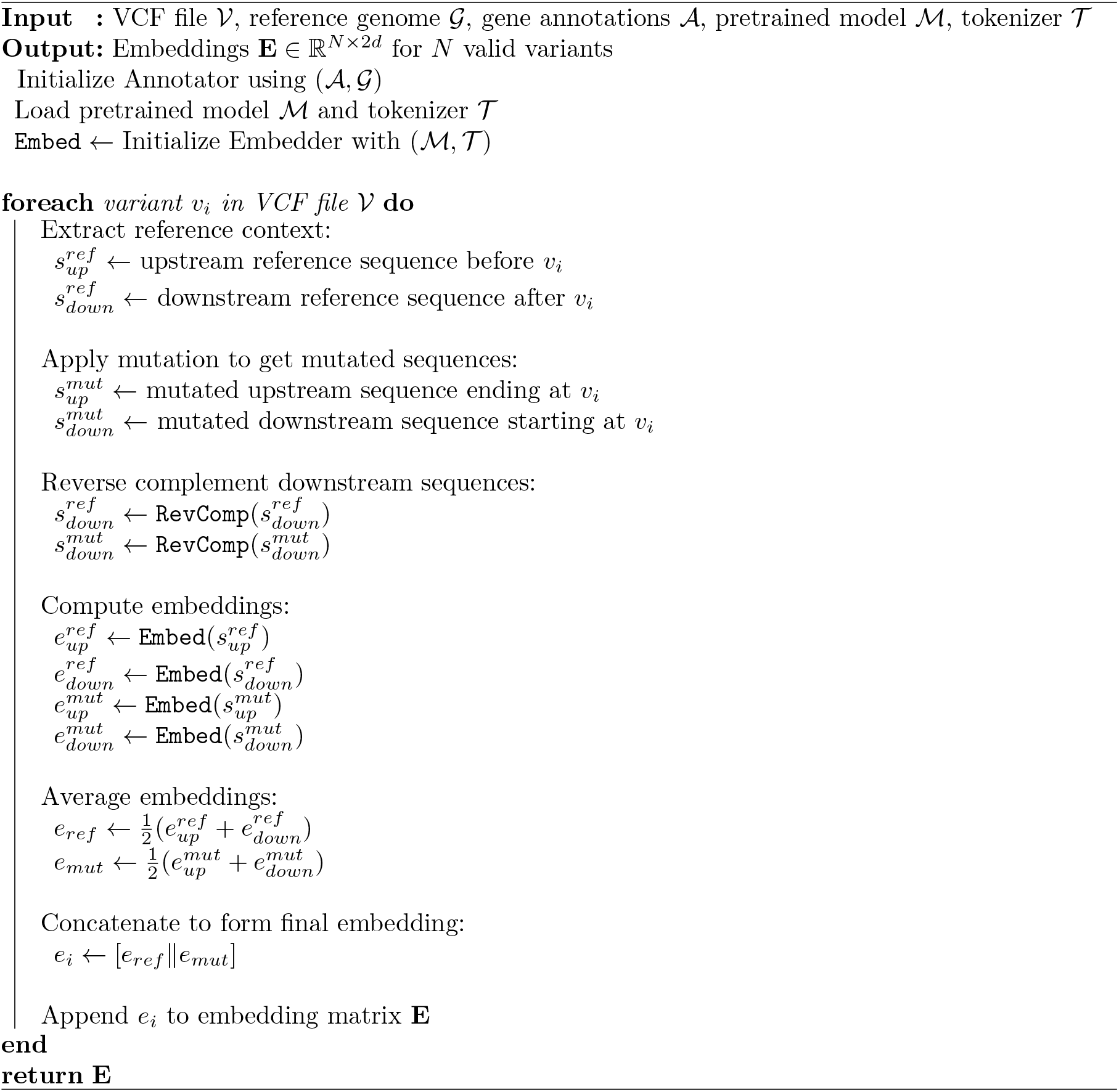

#### A.5 Statistical Tests

We performed a two-sided Wilcoxon signed-rank test (exact method; SciPy library [56]) to evaluate whether the differences in AUC between BioFM and Enformer across 11 cross-validation folds were symmetrically distributed around zero. This non-parametric test does not require normality but assumes symmetry of the differences around the median. BioFM significantly outperformed Enformer (Wilcoxon signed-rank statistic T=1, and p=0.002).

